# Physiological, Biochemical and Microstructural Changes in Rice (*Oryza sativa* L.) under Varying pH Levels

**DOI:** 10.1101/2020.08.17.253914

**Authors:** Ekta, Anil Kumar Singh, Dev Mani Pandey

## Abstract

Soil acidification exerts detrimental effects on rice plant leading to severe reduction in its yield. In the present study, we investigated the physiological, biochemical and microstructural changes in the leaves of rice cultivars, namely, Jhilli Dhan (JD) and Gora Dhan (GD), under varying pH conditions (pH 6.5, 5.5, 4.5 and 3.5). Seedlings were grown at varying pH levels for 14 days under controlled conditions. Root and shoot growth and chlorophyll content were found to be decreased with increasing acidity, whereas electrolyte leakage was increased with increasing acidity. Overall, seedling growth was significantly reduced at pH 3.5, while, it was maximum at pH 6.5 treatment, which might be the result of cumulative altered physiological parameters. Further, biochemical parameters, namely total soluble sugar (TSS), proline content and lipid peroxidation were found to be positively correlated with acidity. Microstructural changes were observed through Swept-Source Optical Coherence Tomography (SS-OCT) and Field Emission Scanning Electron Microscope (FESEM). The thickness between different layers of leaves was found to be disintegrating at low pH conditions and the thickness of parenchyma cells was reduced significantly. FESEM analysis revealed changes in characteristics of stomata under acidic stress. The understanding of physiological, biochemical and microstructural changes in rice leaves under varying pH conditions may help in developing rice with improved tolerance towards soil acidity stress.

## Introduction

About 30% of the world’s land is acidic and around 12% of this acidic land is used for cultivation of crops. Soils having pH < 5.5 in the surface are acid soils which limit the plant growth and yield [1]. Acidic soil leads to poor crop growth, which leads to reduced productivity [2]. A lot of agricultural and industrial activities result in an increased acidity of soil. Improper application of chemical fertilizers, organic matter decay and acid rain are the major factors that increase acidity of the soil [3]. It was also reported that overuse of N fertilizer and intensive farming in China contributes to the acidification of soils [4].

Plant cell maintains cytoplasmic pH in the range of 7.0-7.5 to perform normal functions [3]. But, when the pH falls beyond the optimum level, it creates a negative impact on intracellular pH, disturbing the physiological functions, which are mainly dependent on pH. Decrease in soil pH by 1 unit in the external media, reduces the cytoplasmic pH in root cells by 0.1 units in plants [5, 6]. Plants exposed to strong acidic conditions have enormous impact on root cell structures and its functions [7, 8]. At pH 4.0, legume *Lotus corniculatus* suffered significant damage in the epidermal and cortical cells of roots and depolarized membranes [9, 10]. Reduced root elongations, swollen root hairs and cracks between root meristem region and damaged plasma membranes were observed in arabidopsis [11], alfalfa [12], spinach [13], common bean [14], and barley [15] under acidic stress conditions.

Low pH affects the uptake of water in plants. For example, in case of paper birch (*Betula papyrifera*) grown under low pH conditions, it was found that the water flow rate and hydraulic conductivity were reduced [16]. Low pH conditions decreased the water content and chlorophyll level in *Eucalyptus* leaves [17]. Also, acidic stress leads to reduced root length in plants [18]. It has been reported that soil acidity leads to decreased germination percentage, reduced yield, changed reproductive behaviour and reduced ion transport [19, 20]. The physiological responses of wheat (*Triticum aestivum* L.) cultivars were studied under four different pH levels (3.5, 4.5, 5.5, and 6.5). It was observed that acidic stress reduced growth, biomass, relative water content, and chlorophyll content in all the cultivars [21]. Effect of low pH was investigated in *Citrus sinensis* and *Citrus grandis* at different pH conditions (2.5, 3, 4, 5, or 6) for 9 months and various insights into the causes of low pH-induced inhibition of seedling growth were reported [22]. It was found that pH 2.5 altered the chlorophyll content as well as the water uptake in both species.

The H^+^ ions in acid rain associate with soil cations and displace them from original binding sites; decrease cation exchange efficiency and increase H^+^ concentrations in soil water, which leads to leaching [23]. A high concentration of H^+^ ions in acid soil is also toxic to higher plants, a feature which has been underestimated for several decades [24]. Some basic materials for balancing soil acidity are also lost, as crops are harvested and removed from fields, thus contributing to increased soil acidity. In addition, nutrients such as potassium (K), phosphorus (P), magnesium (Mg), calcium (Ca) and molybdenum (Mo), which are essential for healthy crop growth, become limited to the plants in acidic conditions [25, 26]. The rise in acidic soil is exerting enormous pressure on the world to feed the ever-growing population, which is expected to reach 10 billion by 2050 [27].

Aluminium, the third most abundant element in the earth’s crust, is one of the most toxic elements [28]. In soil with pH 6.0 and above, aluminium forms non-soluble chemical components with only a small proportion remaining in soluble form in the rhizosphere. When the pH of soil reduces below 6.0, Al becomes soluble and causes deleterious effects. Acidity toxicity and Al toxicity cannot be separated since Al is only soluble in acid solution. The aluminium ions (Al^3+)^ cause severe damage to plants by inhibiting root growth and absorption of water and minerals. Al tolerance differs within species among the genotypes [29, 30]. Higher plants employ external mechanisms for adapting to acidic environments. External mechanisms refer to external structures of the root, such as cell wall, cell membrane or chemical exudates including organic acids [31], phenolic compounds [32] and phosphates [33], which prevents Al from entering and accumulating in cells. For e.g., in wheat, tolerance is associated with citrate [34] and malate exudation [35]. Exudation of citrate is an important mechanism for tolerance of *Cassia tora* L. [36], snap bean (*Phaseolus vulgaris* L.) [37], barley [38], and soybean (*Glycine max* L.) [39]. Exudation of oxalate in buckwheat (*Fagopyrum esculentum* M.) [40] and taro (*Colocasia esculenta* (L.) Schott) has been reported [41]. The exudates of organic acid chelate Al^3+^ ions and form non-toxic complexes to avoid interaction of Al with root apices [42].

The higher accumulation of Pro (proline) under acidic stress is linked to upregulated biosynthesis with a decrease in oxidation of Pro [43]. The ROS molecules can oxidize important cellular components including, proteins, lipids, nucleic acids, etc., leading to oxidative damage and destruction of cellular organelles [44], which is indicated by higher levels of malondialdehyde (MDA) in *H. vulgare* [15] and *O. sativa* [45].

Optical coherence tomography (OCT) is a useful imaging method that generates 3D cross-sectional high-resolution images [46]. It is a non-damaging and non-invasive imaging technique and has wide application in the area of ophthalmology, retinal diseases and biomedicine [47, 48]. The application of OCT has been studied in the agriculture sector also. It has been used to observe the morphological differences in different parts of the plant [49, 50]. *In-vivo* studies of fungal infections in leaves of *Capsicum annuum* and its growth were also carried out with the help of OCT [51]. It was found that the internal layers of the disease-affected parts of the leaf had merged into a single thick layer. Similarly, morphological changes in infected apple leaves were also studied [52].With the help of SS-OCT, several distinctive features were found in subsurface boundary regions of treated and healthy leaves in apple trees affected by *Marssonina coronaria*. SS-OCT has been utilized in the identification of the buildup and dissemination of leaf rust disease in wheat leaves that is caused by *Puccinia triticina* [53]. It was reported that proliferation of the fungal hypha leads to the degeneration of parenchyma cells and subsequently the other cell layers too. SD-OCT (spectral domain optical coherence tomography) has been used for the *in-vivo* senescing leaf microstructural changes in *Acer serrulatum Hayata* [54]. Distinctive features among different layers of the leaves were reported; merging of upper epidermis and palisade layers formed thicker layers in red leaves compared to green leaves in *Hayata*. Using the low coherence interferometry method, SS-OCT has been helpful in a non-invasive investigation to characterize the morphological changes of rice leaves [55]. Thus, OCT has become an important tool in the field of agriculture and plant science. OCT is superior to other optical techniques for studying microstructural changes in leaves. Whereas, other optical techniques are invasive and time-consuming, OCT is a fast and robust technique to examine changes in the internal structure of plants. It provides distinguishing features between the healthy and unhealthy leaves.

Rice is the staple food of more than half of the world population and is grown over vast areas of land. However, rice crop is adversely affected in the acidic soil conditions that ultimately lead to reduction in yield of rice. Jharkhand, the 6^th^ largest tribal state of India, is heavily dependent upon agriculture for sustaining its population. However, 90% of the soil in Jharkhand on which agriculture is practiced is acidic with pH<5.5. The state suffers a considerable loss in terms of crop productivity every year due to the acidic soil. The major area of the state is covered with sandy loam to loam soil having pH 4.5-6.5, which has low fertility. The water holding capacity of the soil in the state is very low due to the porous nature of the soil and its undulating topography [56]. Therefore, there is an urgent need to study the effect of low pH on rice plant at multiple levels using modern techniques to increase rice productivity in areas suffering with low pH soil. This study has investigated an array of changes that take place under acidic stress condition in two traditional cultivars of rice that are grown in Jharkhand, namely, JD and GD. Physiological changes were measured through parameters of root and shoot growth, chlorophyll content and electrolyte leakage. Biochemical changes were identified through standard tests, namely, Total Soluble Sugar (TSS), Proline content and Lipid peroxidation. Till now there is no such report, which has utilized SS-OCT to study effect of acidic stress in rice leaves. In this study, SS-OCT based approach has been used to obtain 3D cross-sectional images of rice leaves and to identify the microstructural changes in the cellular region of leaves under varying pH (at pH 6.5, 5.5, 4.5 and 3.5) conditions. FESEM analysis was conducted to determine morphological pattern for varying degrees of acidic stress. The present study reports the impact of acidic stress on rice plant at multiple levels, that is, physiological, biochemical and microstructural. This study is an attempt to gain a holistic understanding of effects of acidic stress on rice leaves. Therefore, it will prove to be of extreme importance in the field of agriculture in general and rice production in particular.

## Material and Methods

### Plant materials and growth conditions

In the present study, rice seeds (*Oryza sativa* L.) of Jhilli Dhan (JD) and Gora Dhan (GD), which are frequently cultivated by the tribal farmers of Jharkhand, India, were obtained from Krishi Vigyan Kendra, Ranchi, Jharkhand, India. Seeds were surface sterilized with 0.1% of mercuric chloride for 1-2 mins and then washed thrice with sterile distilled water, followed by 70% ethanol (for 30s) and again washed three times with sterile distilled water. An equal number of sterilized seeds (30) for both the cultivars (JD and GD) were then soaked on filter paper and kept in Petri plates (9 cm) for 36 h in the dark for germination. Germinated seeds were sown (25 each) on the soil bed in sterile plastic pots. They were further cultivated under controlled conditions and kept in the growth chamber (light, 350 μmol photon m^−2^s ^−1^; temperature, 25 ± 2 °C; relative humidity, 65–70 %) (Saveer, USA) in Department of Bio-Engineering, BIT Mesra, Ranchi, India.

### Acidic stress imposition

Soil samples used in the study were collected from a paddy field, located outside the campus of BIT Mesra, Ranchi, India (23.4123° N, 85.4399° E). The initial pH of the soil was found to be 5.5-5.6. The soil was made acidic by adding water of respective pH 3.5, 4.5 and 5.5 adjusted with 1M HCl in separate pots. About 100ml of water for pH 4.5 and 200ml of water for pH 3.5 was added to decrease the soil pH from 5.5 to 4.5 and 3.5 respectively. Also, about 100ml of water adjusted with 1M NaOH was added to increase soil pH to 6.5 from 5.5. Regular watering was done with the respective pH water to maintain acidity. In order to ensure that the soil pH was maintained, it was checked on regular basis for its respective pH. Germinated seedlings were grown in soil with different pH levels (6.5, 5.5, 4.5 and 3.5) for two weeks (14 days).

### Root and shoot length measurement

Root and shoot lengths of plants of both the varieties were measured after 14d of treatment with various pH.

### Chlorophyll estimation

For chlorophyll estimation small pieces of plant leaves were cut and homogenized with 1ml of 80% acetone using mortar and pestle [57] at room temperature. The homogenate was further centrifuged at 5000 x g for 10 min. The absorbance of chlorophyll containing acetone extract was taken immediately using spectrophotometer (Perkin Elmer, USA, Lambda-25) at 663 and 645 nm, using 80% acetone as a blank. Following formulas were used to calculate the Chl a, Chl b and total Chl contents.

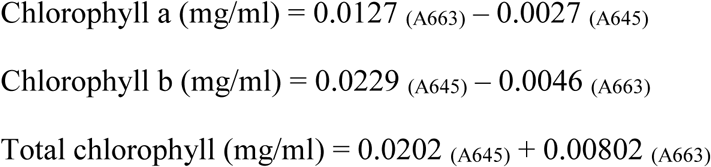

### Electrolyte leakage determination

Electrolyte leakage (EL) was estimated using young leaf tissues [58]. Total 1g of plant sample washed with distilled water was placed in test tubes having 15ml of distilled water and incubated for 12 hours at 24°C. The first electrolytic conductivity (EC1) was measured with the help of conductivity meter. Further, the same sample was autoclaved for 20 min at 120°C and the second electrolytic conductivity (EC2) was measured. The electrolyte leakage was calculated as:

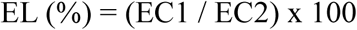

### Total soluble sugar

About 100 mg of sample was homogenized in 3 ml of 80% methanol and was incubated at 70°C for 30 min. After incubation, equal volume (500 µl) of extract and 5% phenol was mixed with 1.5 ml of 95% H_2_ SO_4_ and further incubated in dark for 15–20 min. Absorbance was measured in spectrophotometer at wavelength of 490nm [59].

### Proline estimation

About 0.5g of leaf sample was homogenized in 10ml of (3%) sulfosalicylic acid and further centrifuged at 12,000 rpm at room temperature [60]. About 2ml of supernatant was mixed with same volume of acid ninhydrin and glacial acetic acid and incubated for 1 h at 100°C. The reaction was terminated in an ice bath, and 1ml of toluene was added. The absorbance of toluene phase was measured at 520nm.

### Lipid peroxidation

The level of lipid peroxidation (LP) in control and treated tissues was determined by measuring malondialdehyde (MDA) content via 2-thiobarbituricacid (TBA) reaction [61]. About 100 mg of leaf tissue was homogenized in 500 µl of 0.1% (w/v) TCA and centrifuged for 10 min at 13,000 g at 4°C. Further 500 µl of supernatant was then mixed with 1.5 ml 0.5% TBA and incubated in water bath at 95°C for 25 min. Mixture was then incubated on ice for 5 min for termination of reaction. Absorbance of mixture was measured at 532 and 600 nm.

### Leaf sample preparation for OCT imaging

For OCT imaging, fresh leaves were used during day time. The pots containing plants were brought to the SS-OCT system after 14 days of acidic stress treatment. Leaf samples within the plant were hydrated with sterile distilled water. Firstly, the leaf was washed with distilled water and then a small portion of leaf was mounted on the glass slide for SS-OCT imaging. It was ensured that the images of fresh leaves were taken and damaged leaves were avoided.

### SS-OCT system instrumentation

The leaf samples were processed using SS-OCT (Fig. 1) setup. SS-OCT encodes physiological features of plant through optical interferometry in form of A scans and B scans. The image acquired through SS-OCT was processed using an algorithm in MATLAB to enhance the layers by averaging and attenuating signal to noise (S: N) ratio. The B-scans are processed to get A-line scans for different sections.

**Fig. 1.**
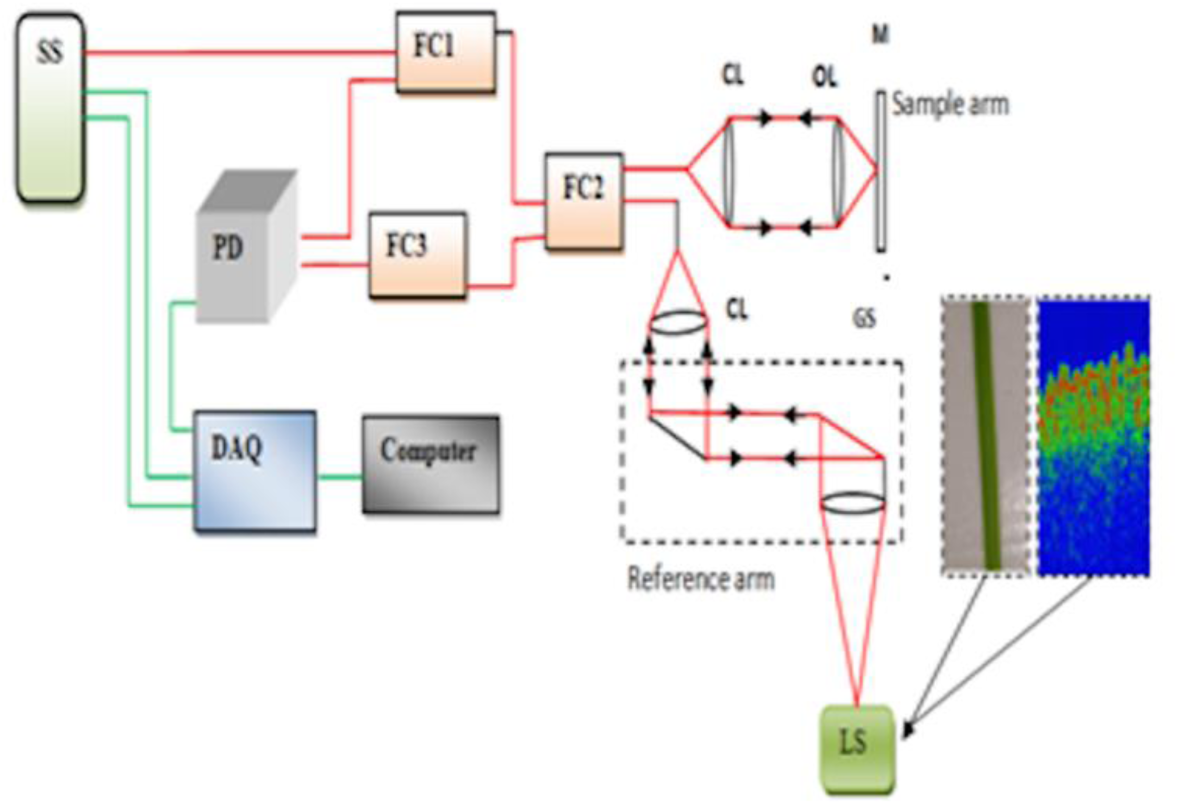
Schematic representation of laser swept source OCT; FC1- 50:50 fiber coupler 1; FC2- 50:50 fiber coupler 2; FC3- 50:50fiber coupler 3; CL- Collimating Lens; OL- Objective Lens; DAQ-Data Acquisition; GS- Galvo-scanner (Cambridge Technology dual-axis); LS- Leaf Sample; DAQ- Analog to digital conversion data acquisition board and PD- Photodetector.

The protocol used for scanning was: Pixels per A-scan: 2048, No. of A-scans per B-scan: 100, Scanning distance: 2mm, Time per B-scan: 1 ms, Axial imaging resolution: ∼5 µm, Transverse imaging resolution: ∼14 µm, Axial imaging range: 3.2 mm [53].

### Determination of scattering coefficient (µs) from OCT data

The scattering coefficient has been calculated for the given rice OCT data using the Beer-Lambert’s law. The equation used for calculating the scattering coefficient is [62]:

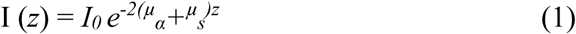

Here,

I_0_ = incident light

μ_a_ = absorption coefficient

μ_s_ = scattering coefficient

z = depth from the surface of the plant tissue

The absorption coefficient of most of the plant tissues is very low in NIR (near infrared region) region, hence we can neglect coefficient of absorption (µ_a_) in equation 1, which gives;

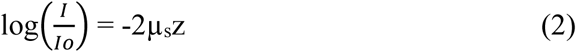

For the determination of variations in the rice seedlings under different pH conditions, an average of 50 pixels of 100 A-scans, were computed at five different regions of the image.

### FESEM

A field emission scanning electron microscope (FESEM, ZEISS Sigma 300) was used to determine leaf surface characteristics as well as the stomata characteristics [63] under extreme acidic condition (pH 3.5) and control conditions (pH 6.5) for both the rice varieties [64] at 1.00 K X and 5.00 K X magnification. The leave samples of both the rice varieties were fixed in 2.5% glutaraldehyde and incubated at 4°C overnight. The dehydration of cells was carried out using a different concentration of ethanol and then subjected to air-drying.

### Statistical analysis

Each experiment was conducted with three biological and three technical replicates, which are expressed as mean with standard deviation (mean ± SD). ANOVA (analysis of variance) was performed and compared among the means using the Bonferroni multiple range test at P < 0.05. The results were transformed to graphical representation using Graph Pad Prism software (version 8.3.1).

## Results

### Effect of pH on seedling growth

After 14 days of acidic stress imposition, both the rice varieties showed similar growth pattern under different pH conditions. In case of both Jhilli Dhan (JD) and Gora Dhan (GD) varieties, the maximum seedling growth was observed at pH 6.5 (control). In JD, the shoot length was slightly reduced by 10.21% and 25.98% at pH 5.5 and 4.5 respectively, when compared with control (pH 6.5), while at pH 3.5, shoot length was drastically reduced by 55.55%. However, in GD, shoot length decreased by 21.4%, 36.15% and 50.07% at pH5.5, 4.5 and 3.5 respectively when compared with control plants (pH 6.5). Root length was found to be reduced by 35.3%, 60.2% and 65.7% in GD variety while 23.8%, 42.7% and 61.7% in case of JD at pH 5.5, 4.5 and 3.5 respectively, with respect to pH 6.5. The pH value had a direct relation with root length and shoot length which implies that at low pH, root length and shoot length were found to be significantly reduced (Fig. 2A, B).

**Fig. 2.**
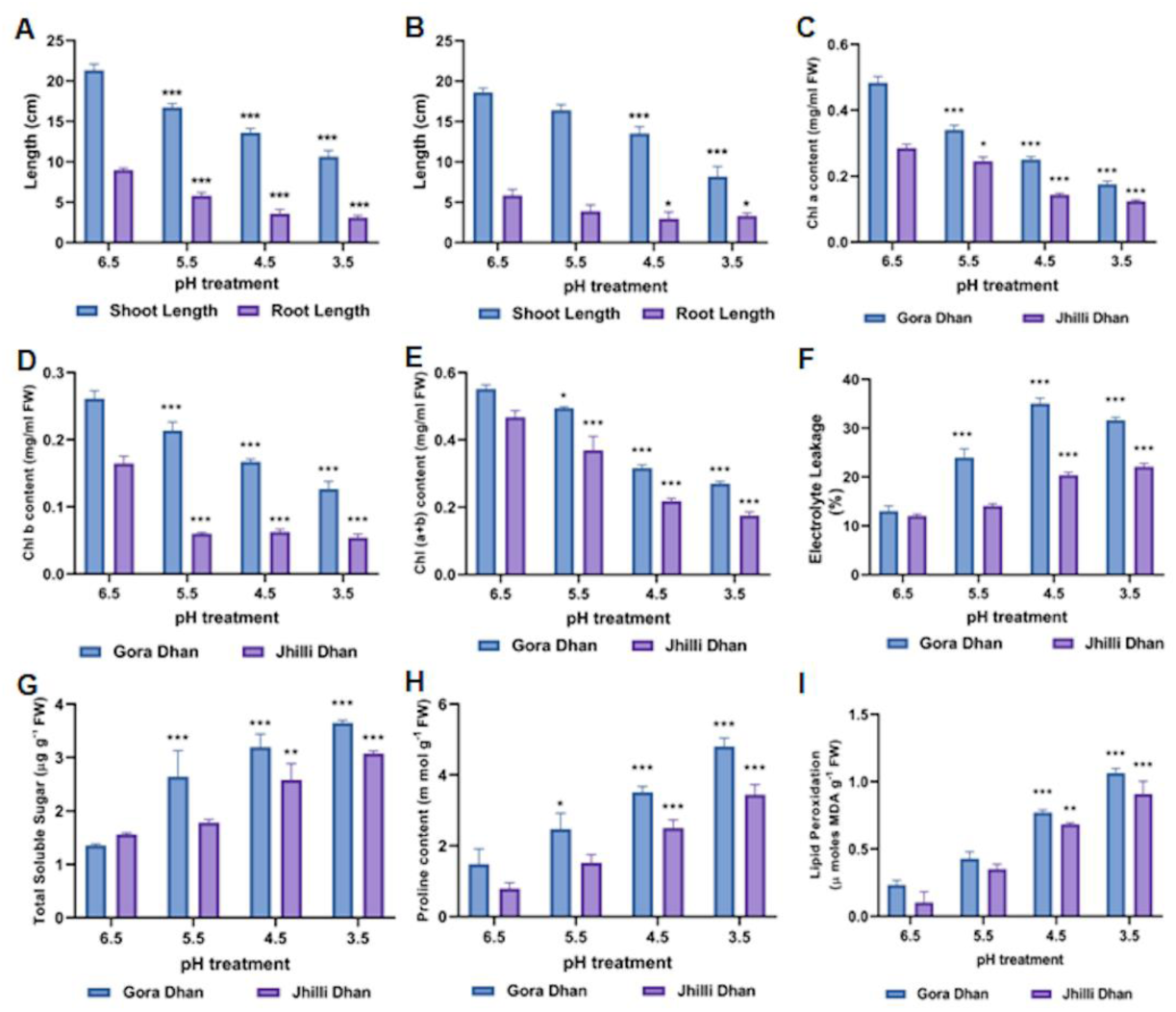
Physiological and biochemical response of low pH on rice seedlings. (A) and (B) represent the root and shoot length measured after 14 d of acidic stress treatment in GD and JD, respectively. (C), (D), (E) represent the effect of low pH on chl a, chl b and total chl contents, respectively in the rice seedlings, (F) electrolyte leakage, (G) total soluble sugar, (H) proline content, (I) lipid peroxidation of GD and JD varieties. Bars represent standard deviation among three biological replicates and asterisk ‘*’, ‘**’, ‘***’ represent significant difference at p < 0.05.

### Effect of pH on chlorophyll content and electrolyte leakage

Acidic stress was found to be detrimental to the photosynthetic pigments in both the rice varieties. Both JD and GD varieties exhibited the reduced Chl a content at low pH level and in a similar fashion Chl b content was also found to be reduced. Maximum reduction in Chla and Chl b content was observed at pH 3.5. Total Chl content was also recorded at its lowest level at pH 3.5 in case of both JD and GD. Total Chl was found to be significantly reduced for both the genotypes, with values being lower by 21.08%, 53.5% and 62.54% in JD and 10.43%, 42.7% and 50.8% in GD at pH 5.5, 4.5 and 3.5, respectively, when compared with the control, that is, pH 6.5(Fig. 2C-E).

Leaf electrolyte leakage (EL) was found to be increased as the pH decreased from 6.5 to 3.5 (Fig. 2F). A significant increase in leaf EL was observed in plants treated with pH 5.5, 4.5 and 3.5 when compared with the control. In JD, the EL content was found to be increased by 14.5%, 40.9% and 45.5% at pH 5.5, 4.5 and 3.5, respectively when compared to pH 6.5. In GD, the EL content was found to be increased by 45.9%, 62.9% and 58.9% at pH 5.5, 4.5 and 3.5 respectively when compared with pH 6.5.

### Effect of low pH on total soluble sugar, proline and lipid peroxidation

An increase in acidity with respect to pH6.5 was followed by significant increase in accumulation of soluble sugar. In comparison to control condition (pH6.5), progressive increment in TSS content was recorded in samples under acidic stress at pH 5.5, 4.5 and 3.5, respectively (Fig 2. G). The quantitative increase in TSS content was found to be 94.9%, 135.8% and 169.3% at pH 5.5, 4.5 and 3.5, respectively, when compared to pH 6.5 for GD variety. In a similar fashion the increase recorded in case of JD was 15.9%, 80.7% and 92.3% at pH 5.5, 4.5 and 3.5 respectively, with respect to pH6.5. The maximum percentage increase in TSS content (∼170 %) was recorded at pH 3.5 in case of GD variety. For both the varieties the degree of accumulation of TSS was found to be least at control condition (pH 6.5) in comparison to treated condition (pH 5.5, 4.5 and 3.5).

Proline accumulation at varying acidity levels (pH 6.5, 5.5, 4.5 and 3.5) was recorded for JD and GD variety. It was observed that the proline content under different levels of acidity increased in a dose dependent manner. The quantitative increase in proline content for JD variety was 91.9%, 217.7% and 335.8% at pH 5.5, 4.5 and 3.5, respectively in comparison to pH 6.5. Similarly, the percentage increase in proline content for GD variety was 67.1%, 137.6% and 224.3% at pH 5.5, 4.5 and 3.5, respectively in comparison to pH 6.5. The highest increase in proline content (∼336%) was recorded at pH 3.5 for JD variety when compared to control (pH6.5). In addition, for every level of acidic stress the proline content was found to be higher in JD variety in comparison to GD variety (Fig. 2H).

Lipid peroxidation test was performed to check the integrity of membrane at varying levels of acidity. Lipid peroxidation (measured as MDA) in the rice leaf tissue was found to increase in a progressive manner in correspondence with increasing acidity level. The quantitative increase in MDA content for JD variety was 244.9%, 571.8% and 794% at pH 5.5, 4.5 and 3.5, respectively in comparison to pH 6.5. In a similar manner the percentage increase in MDA content for GD variety was 77.5%, 234% and 372.8% at pH 5.5, 4.5 and 3.5, respectively, in comparison to pH 6.5. The highest increase in MDA content (794%) was recorded at pH 3.5 for JD variety as compared to control (pH6.5) (Fig 2. I). In addition, for every level of acidic stress, the MDA content was found to be higher in JD variety in comparison to GD variety. Further, at control condition (pH 6.5), MDA content was least compared with the rest of the treatments for both the varieties.

### Effect of acidity on rice leaf: SS-OCT Analysis

In our study, rice leaves were observed at varying pH of 6.5 (control), 5.5, 4.5 and 3.5. The Fig. 3A shows a white dotted line within the region of interest indicating the scanned position of the leaf specimen while, Fig. 3B represents the microstructural cellular formations, such as the waxy layer of upper cuticle, thick epidermal cell layer, aerenchyma cellular regions, oval shaped parenchyma cells, vascular bundles and stomata. In OCT images (A-scans and B-scans), the thickness between different layers of treated and control leaf samples were observed. A-scan is related to longitudinal scan and gives us a single dimensional image (Z-axis). B-scan is related to transverse scan and gives us a two dimensional image (XZ or YZ axis). The analysis of A-scans and B-scans led us to the conclusion that higher acidity has a negative effect on growth of plants as it reduces the rate of photosynthesis in leaves by altering its structure, specially the parenchyma cell layer. The detailed analysis is described in the subsequent paragraphs.

**Fig. 3.**
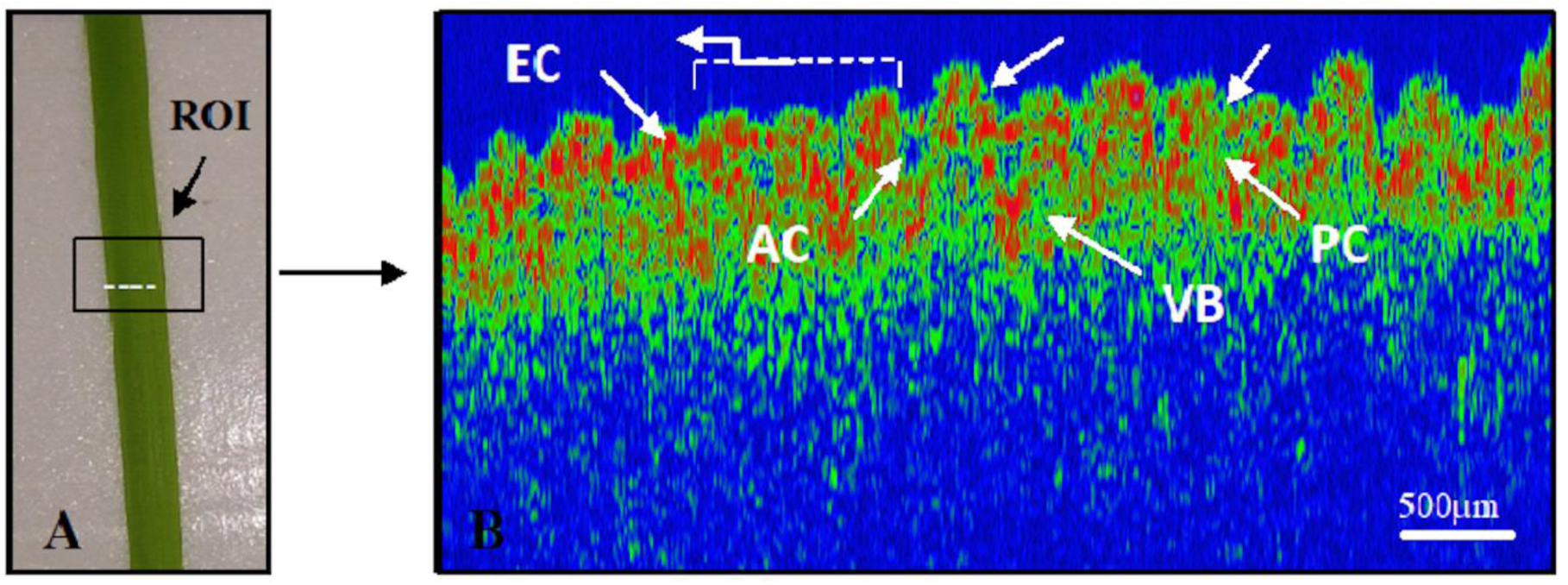
Two-dimensional (2D) SS-OCT image. (A) Treated acidic rice leaf; black box represents the region of interest ‘ROI’ (B) real time, *in-vivo* OCT B-scan. UC: Upper Cuticle, V: Veins, AC: Aerenchyma Cell, PC: Parenchyma Cell, VB: Vascular Bundle, EC: Epidermal cell and ST: Stomata.

### B-Scan Images

The leaf specimens were observed and imaged on an adaxial surface. Figure 4, represents the B-scan SS-OCT cross-sectional depth images of GD and JD varieties. The constant scan range for all the specimens under different pH conditions was kept constant at 2mm for both the varieties. The white box on the B-scan OCT image represents the corresponding region of interest. In Fig. (4 A-H), OCT B-scan images show the microstructural variations present under different pH treatments representing the distinct layers of the leaf, i.e., cuticle, epidermal cells, parenchyma cells, vascular bundles, and stomata.

**Fig. 4.**
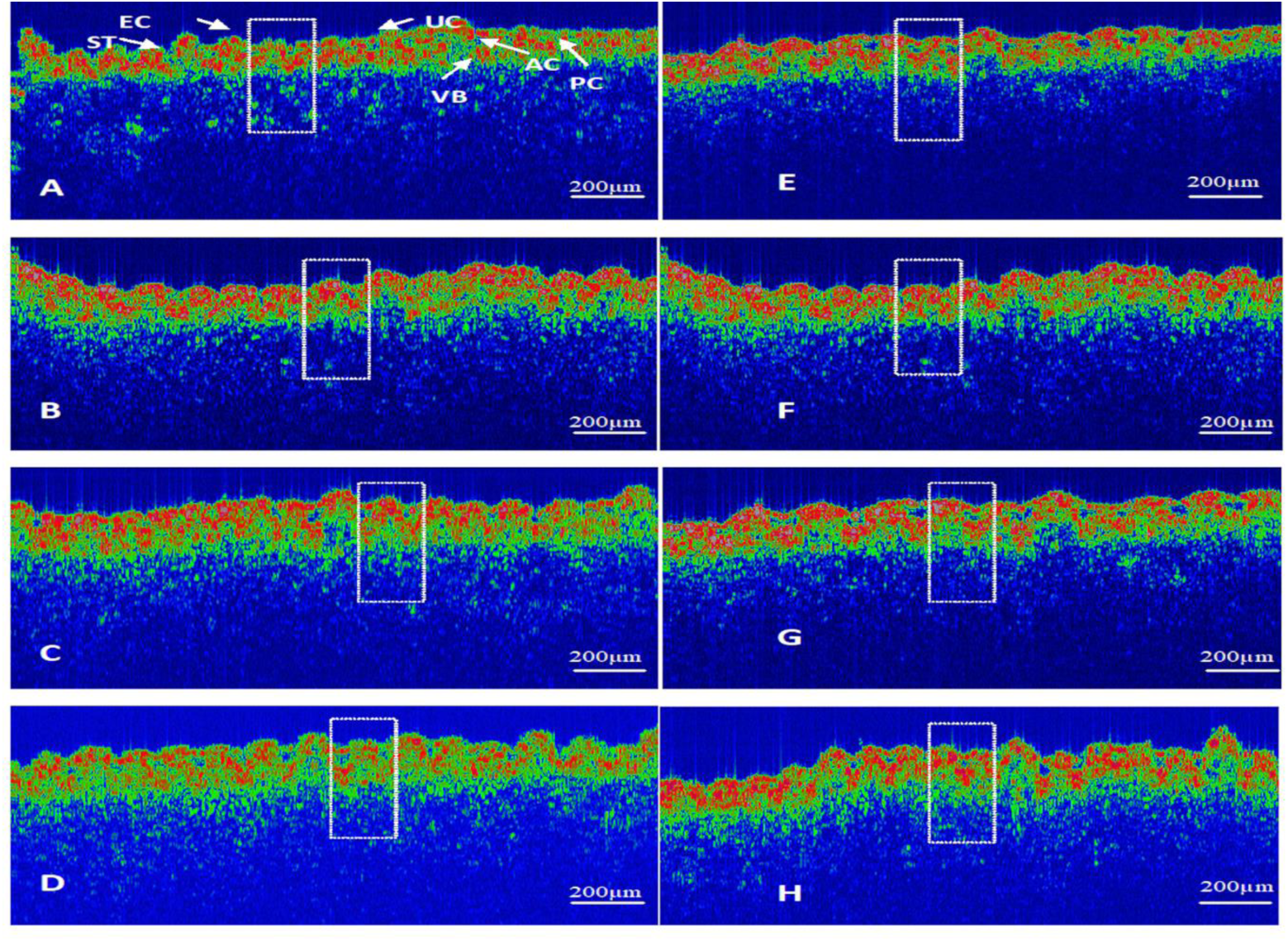
Two-dimensional (2D) SS-OCT image analysis. (A), (B), (C) and (D) represent SS-OCT B-scan images of Gora Dhan at pH 3.5, 4.5, 5.5 and 6.5, respectively. (E), (F), (G) and (H) represent SS-OCT B scan image of Jhilli Dhan at pH 3.5, 4.5, 5.5 and 6.5, respectively. The white rectangular box represents the region of interest (Scale bar: 200µm). UC: upper cuticle, EC: epidermal cell, PC: parenchyma cell, VB: vascular bundles, ST: stomata.

At pH 4.5 and 3.5, the parenchyma cell layer was found to be indistinguishable and merged with the epidermal cell layer leading to a thickened layer (Fig. 4 A, B). While, at pH 5.5 the epidermal cell layer has thickened, as the parenchyma cells have started to merge with upper layer due to acidic stress (Fig. 4C). However, at pH 6.5, the B-scan image of GD showed different layers of the leaf which indicates that the leaf is in healthy condition and all the three layers are visible (Fig. 4D).

Similar observations were recorded in case of JD, which are shown in Fig. 4E-H. The increase in acidity from pH 6.5 to pH 3.5 was accompanied by disintegration of parenchyma cell layer and its merging with upper layer leading to a thickened layer at higher acidity distinguished from three distinct layers at lower acidity.

The above observations in case of both JD and GD varieties lead us to the conclusion that soil acidity leads to disintegration of parenchyma cell layer as it merges with epidermal cell layer, thereby having an adverse effect on photosynthetic pigments. Hence, higher acidity reduces the photosynthesis in leaves leading to reduced growth of plants.

### A-Scan Images

In order to obtain the A-scan depth profile peaks, a program was coded to search for the intensity of depth peaks. These peaks were used to analyse and compare the thickness in the layers of leaf (Fig. 5). The thickness between two layers of a leaf is akin to corresponding distance between A-scan profile peaks. The peak intensities of a healthy A-scan are significantly distinguishable and such clear peaks are not obtained under increased acidic conditions. The appearance of clear peaks confirm the morphological layer information of leaf while the disappearance of clear peaks confirm the morphological changes that have occurred in the leaf due to acidic stress treatment. At pH 6.5 (Fig. 5D, H), peaks in A-scan plot showed different layers of the leaves. On the other hand, at pH 3.5 (Fig. 5A, E), the absence of peak for parenchyma cell layer was visible. At pH 5.5 (Fig. 5C, G) and at pH 4.5 (Fig 5B,F), the second order peaks have broadened. This substantiates our inference that with increasing acidity the parenchyma cell layer starts merging with epidermal cell layer and finally merges to form a thickened layer.

**Fig. 5.**
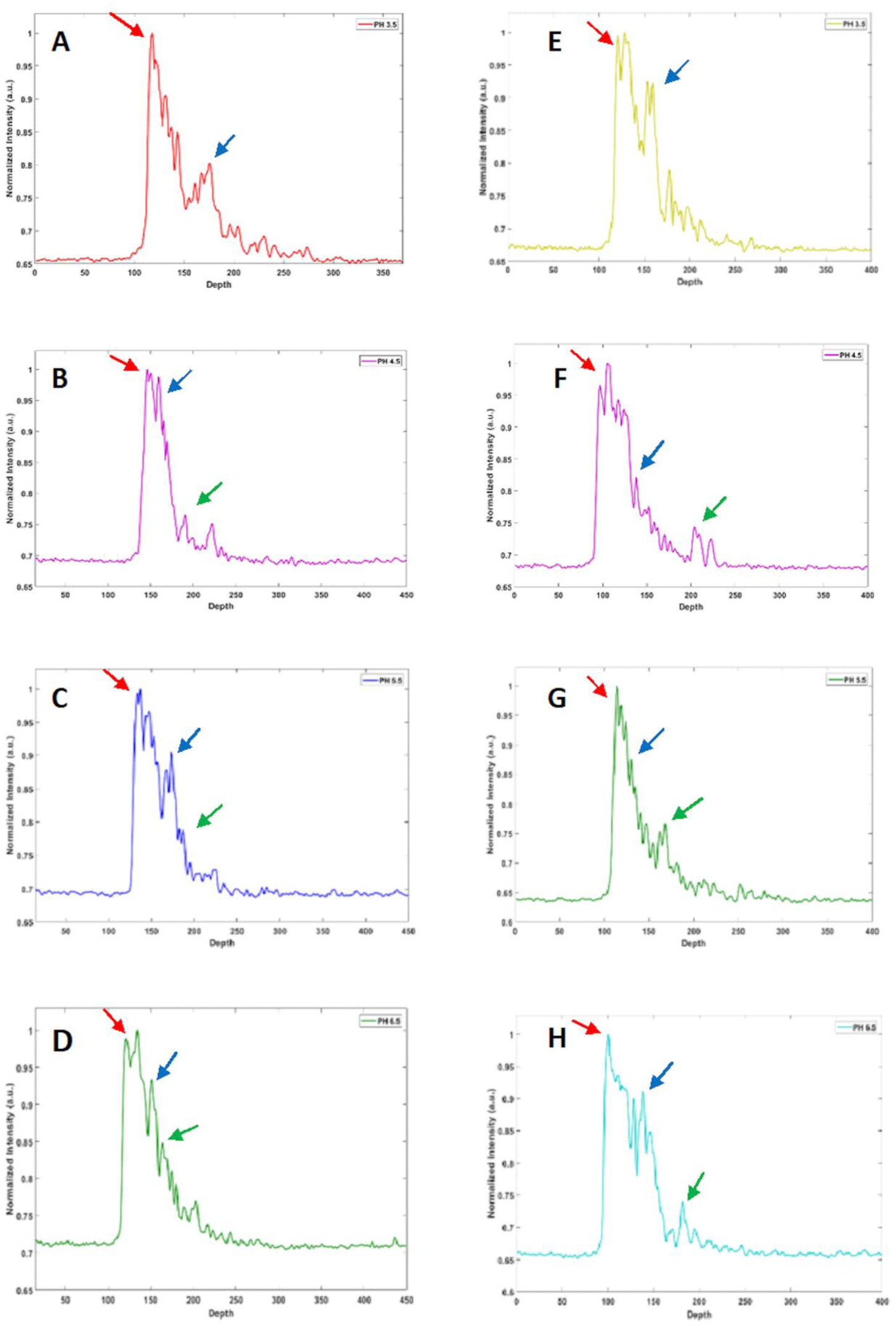
Corresponding A-line depth profile. (A), (B), (C) and (D) represent averaged A-scan of the region of interest for Gora Dhan at pH 3.5, 4.5, 5.5 and 6.5 respectively. (E), (F), (G) and (H) represent averaged A-scan of the region of interest for Jhilli Dhan at pH 3.5, 4.5, 5.5 and 6.5 respectively. Red arrow: upper cuticle; Blue arrow: epidermal cell; Green arrow: parenchyma cell.

In order to analyse the overall effect of acidity on rice plant at different pH levels normalized depth profiles were plotted (Fig. 6 A, B), wherein it was observed that the location of uppermost layer, i.e., the upper cuticle layer is almost fixed (red arrow in Fig. 6). Further, degeneration of the upper epidermis and parenchyma layer was observed with increase in acidity, which was indicated by broadened peak.

**Fig. 6.**
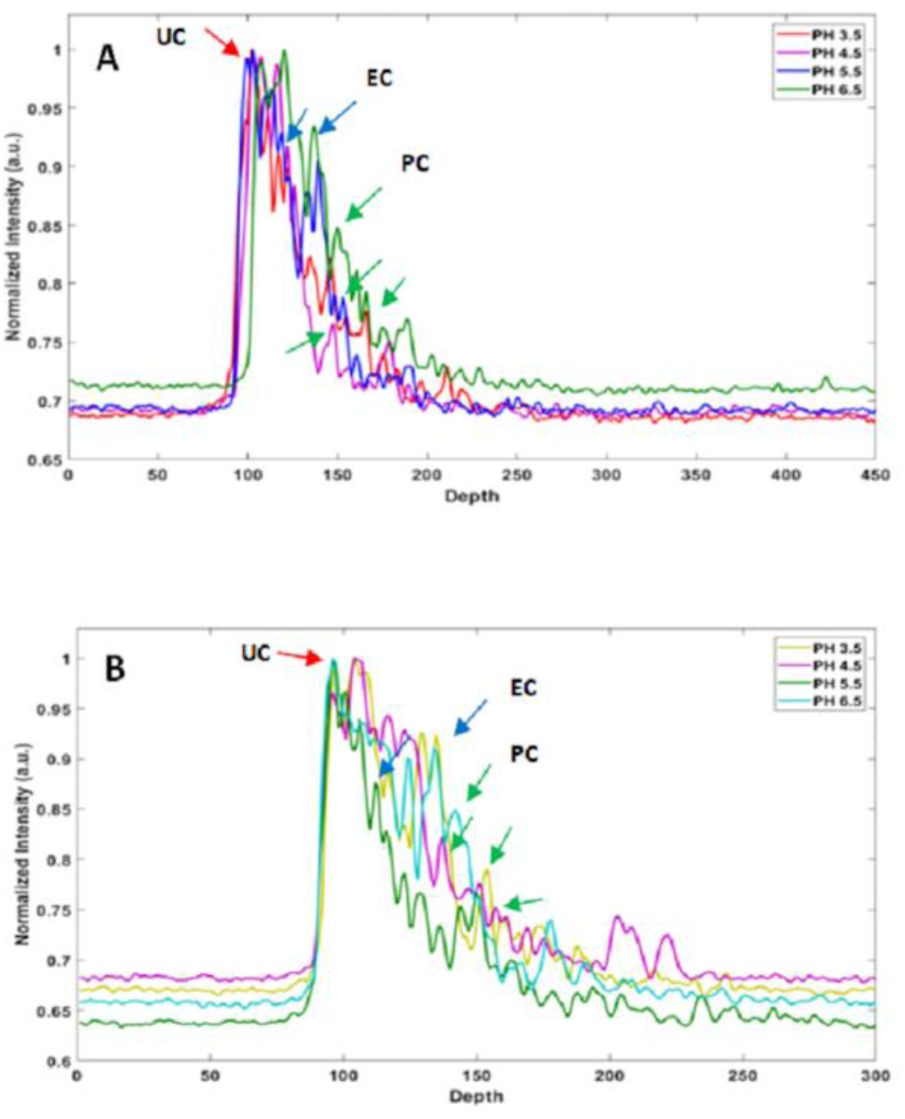
A comparison between depth profiles (averaged A-scans) (A) Cumulative averaged A-scan depth profiles for the region of interest for Gora Dhan. (B) Cumulative averaged A-scan depth profiles for the region of interest for Jhilli Dhan. UC: Upper Cuticle; EC: Epidermal Cell; PC: Parenchyma Cell.

The degeneration of parenchyma cell and its subsequent merging with epidermal cell layer can be considered as one of the manifestation of increased acidity revealed through structural variation in leaves. Thus, it might be useful to identify the leaf layer structural change to understand effect of acidic stress on plant. Effect of acidity on the parenchyma cells and the chlorophyll content showed correlation with different peak intensities obtained in the depth profiles.

### Scattering Coefficient

Further, to quantify the changes in terms of intensity at different pH levels, the scattering coefficient of tissue layers was calculated and plotted (Fig. 7 A, B). Scattering coefficient can be defined as amount of light scattered per unit distance as light travels in a tissue. It is an intrinsic optical property of tissues. The calculated data for scattering coefficient is shown in Table 1. The scattering coefficient (mm^−1^) for JD variety at different pH levels are: 9.76 (pH 3.5), 8.67 (pH 4.5), 8.17 (pH 5.5) and 7.89 (pH 6.5). The scattering coefficient (mm^−1^) for GD variety at different pH levels are: 12.3 (pH 3.5), 10.5 (pH 4.5), 9.43 (pH 5.5) and 8.02 (pH 6.5).The results indicate increasing scattering coefficient with increasing acidity. The value of scattering coefficient peaks at pH 3.5 for both varieties of rice. The higher value of scattering coefficient corresponds to greater scattering of light at same penetration depth.

**Table 1.** Scattering coefficients of Gora Dhan (GD) and Jhilli Dhan (JD) rice leaf tissue under different conditions of pH treatment.

| pH treatment | Scattering Coefficient ($\text{mm}^{-1}$) GD | Scattering Coefficient ($\text{mm}^{-1}$) JD |
| --- | --- | --- |
| pH 3.5 | 12.3 | 9.76 |
| pH 4.5 | 10.5 | 8.67 |
| pH 5.5 | 9.43 | 8.17 |
| pH 6.5 (control) | 8.02 | 7.89 |

**Fig. 7.**
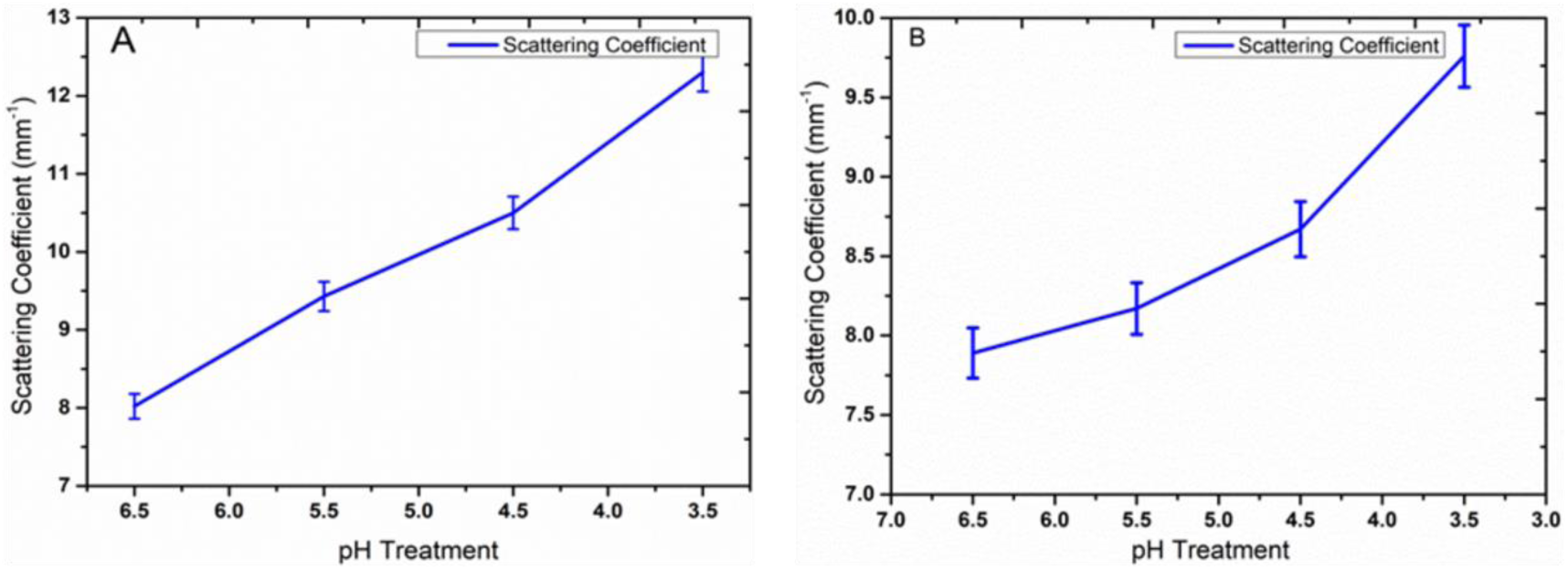
Local scattering coefficient of leaf tissue layers at different pH level. (A) Significant increase in scattering coefficient of Gora Dhan as pH varies from 6.5-3.5. (B) Significant increase in scattering coefficient of Jhilli Dhan as pH varies from 6.5-3.5.

### Effect of low pH on stomata: FESEM analysis

Plant shows morphological changes in response to acidic stress. There are reports that acidic stress has a negative impact on morphological structures of leaf, including stomatal density as well as the opening and closing of stomata. In this study, both JD and GD rice varieties were studied for differences in stomatal characteristics. It was observed that the number of stomata was decreased at lower pH for both the varieties. Figure 8shows reduced number of stomata at pH 3.5 as compared to control. The stomata were found to be open and the pore was clearly observed at pH 6.5 in both, JD and GD rice (Fig 8. E, G). Whereas, in pH 3.5 conditions, stomatal pore length as well width were found to be reduced significantly in case of JD (Fig 8. F), while in case of GD the stomata were almost closed and no pore was visible at pH 3.5 (Fig 8. H). These observations suggested that, higher acidity leads to reduction in stomatal density as well as changes in the structure of stomata. Consequently, it can also be inferred that at higher acidity, photosynthetic rate reduces due to changes in stomatal conductance.

**Fig. 8.**
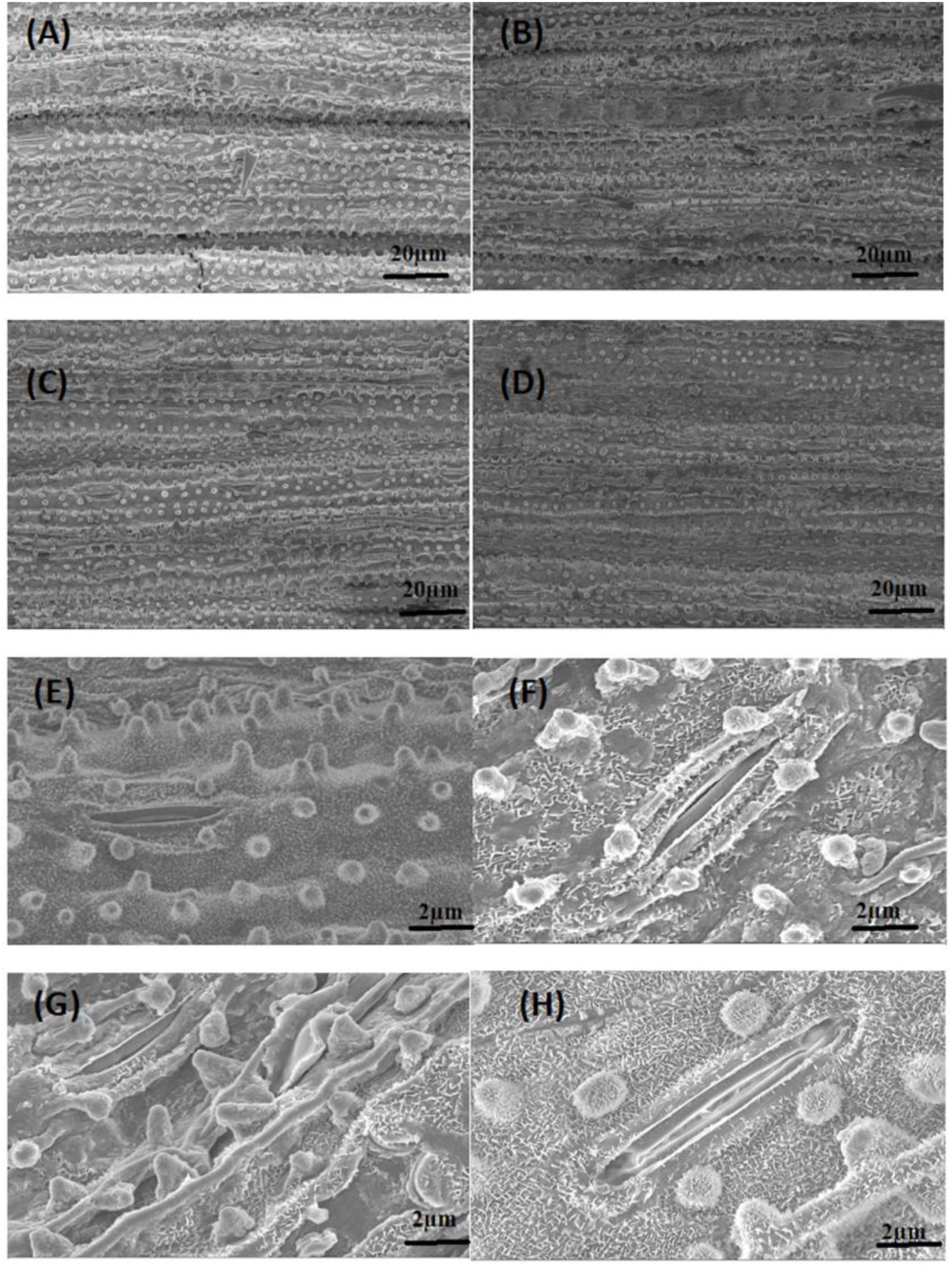
Stomatal frequency of rice after 14 d of acidic stress. (A) Jhilli Dhan at pH 6.5 (B) Jhilli Dhan at pH 3.5 (C) Gora Dhan at pH 6.5 (D) Gora Dhan at pH 3.5; using a field emission scanning electron microscopy (FESEM) at 1.00 K X magnification; scale bar = 20µm. (E) Jhilli Dhan at pH 6.5 (F) Jhilli Dhan at pH 3.5 (G) Gora Dhan at pH 6.5 (H) Gora Dhan at pH 3.5; using a field emission scanning electron microscopy (FESEM) at 5.00 K X magnification; scale bar = 2µm.

## Discussion

### Acidic stress altered the root and shoot length of rice seedlings

Acidic stress adversely effects crop productivity by altering various physiological and biochemical processes. Acidic soils, having pH<=5.5 are major limitations to the crop production area [2]. In the present study, low pH stress (acidity) was found to negatively influence various morpho-physiological parameters of rice seedlings. Exposure of rice seedlings to acidic pH lead to a significant reduction in the seedling growth in terms of root and shoot length in both the rice cultivars. It has been reported that reduction in root elongation is due to the higher H^+^ toxicity [45, 22] and due to reduction in cell division [65]. The effect of low pH has been previously reported on rice seedlings in hydroponic conditions, where seedling growth was affected significantly as the pH decreased in the dose dependent manner (from pH 5.5 to 3.5). The reduction in root length and root surface area was reported under low pH conditions [66]. Effect of low pH conditions (pH 5.5-2.5) on seedling growth was analyzed in *Citrus* and it was reported that low pH conditions greatly reduced the seedling growth as well as altered many physiological parameters, like, seedling growth, concentrations of nutrient elements in root, stem, and leaf, chlorophyll a fluorescence; relative water content, total soluble protein level, H_2_O_2_ production and electrolyte leakage in roots and leaves at pH 2.5 [22].

### Acidity increased electrolyte leakage

Plants being sessile cannot escape the environmental stresses. It has been previously reported that the electrolytes leaked out of the cell in wheat cultivars as the acidity of the soil increased [21]. Similar observations have been reported in the present study, where, EL was found to be maximum at the lowest pH (3.5) condition, indicating that acidic stress leads to damage of membranes in the cell causing the electrolytes to come out of the cell. An increase in EL in *Cucurbita pepo* was also found under pH 3.5 [67]. Similar observations were also reported in case of roots of paper birch (*Betula papyrifer*) seedlings, where low pH (4.0) was found to cause reduction in water flow rate as well as hydraulic conductivity [68].

### Extreme pH conditions damaged the photosynthetic pigment rate

Extreme acidity not only causes reduced uptake of water but also damages the photosynthetic pigments [44]. In the present work, Chl a, Chl b and total Chl contents were found to be decreased with reduction in pH. The maximum and minimum chlorophyll contents were observed at pH 6.5 and pH 3.5, respectively. In *Eucalyptus*, reduced Chl content and aluminium toxicity due to damage in chloroplast envelopes was observed at pH 3.0. Low pH also decreased the relative water content in *Eucalyptus* roots, stems, and leaves [69]. Similarly, in case of wheat cultivars acidic pH stress caused reduction in the Chl (a+b) content as compared to the control [21]. The same was observed in case of *Arabidopsis thaliana*, where the Chl content was found to be reduced due to acidic stress condition [70].

### Acidic stress lead to accumulation of TSS and Proline

Total Soluble Sugar (TSS) is a compatible osmolyte which maintains membrane stability, thereby, helping plants to overcome several environmental stresses. Proline is an osmoprotectant which helps plant withstand stress by assisting in the process of osmoregulation. The accumulation of compatible osmolytes such as proline, soluble sugars, glycine betaine, trehalose etc. help plants to withstand abiotic stresses through maintenance of osmotic turgor [71, 72]. Past studies have highlighted the correlation between increased levels of TSS and proline content and increased stress tolerance in plants [71, 73]. Similarly, our study also reports an increase in TSS and proline content when GD and JD varieties of rice are exposed to acidic stress condition. For both varieties, the TSS and proline content were found to be increased significantly in a progressive manner in correspondence to increasing acidity levels (pH 5.5, 4.5 and 3.5) in comparison to control condition (pH 6.5). The greater increase in level of TSS and proline content for JD variety in comparison to GD variety under acidic stress can be associated with higher level of stress tolerance in JD rice seedlings. A similar study has been reported in case of wheat seedlings, wherein, it was found that as acidity increased from pH 6.5, 5.5, 4.5 and 3.5, proline content increased significantly. The highest proline content was recorded at pH 3.5 for wheat seedlings [21]. Higher accumulation of proline may help to avoid physiological injury and maintain major physiological processes [44]. An elevated proline content under acidity stress is attributed to protection against oxidative injury and maintenance of water status [44]. Hence, higher proline content in JD variety under stress can provide better protection against injuries in comparison to GD variety.

### Acidity enhanced peroxidised lipids under acidic stress

MDA content indicates extent of lipid peroxidation in living tissues. An increase in MDA reflects more membrane damage and *vice-versa*. Our results are consistent with observations of oxidative damage and elevated content of MDA in *H. Vulgare* under acidity stress [15]. Similarly, ROS content was enhanced with increased acidification towards pH 4.5 in *P. sylvestris* [74]. Acidic stress has been found to retard growth in *Arabidopsis* by over-accumulation of H_2_O_2_ and MDA [70]. An increase in MDA was followed by plants suffering from severe oxidative stress resulting in consequent membrane damage. Similar to the findings of this study, MDA content was recorded to be elevated significantly in wheat cultivars as the acidity increased from pH 6.5 to pH 3.5 [21].

### Microstructural variation under low pH condition

As of now, very little information is available about the effects of acidity on rice plants at microstructural level due to lack of research on the same. In the present study, we used SS-OCT and FESEM, to study rice plants at microstructural level under varied pH conditions. Our results demonstrated that a decrease in pH results in reduction in growth of plants that can be explained through alterations in physiological parameters caused due to structural changes.

The main focus of our study was to study the effect of varying pH on microstructural cellular formations, such as the waxy layer of upper cuticle, thick epidermal cell layer, aerenchyma cellular regions, oval shaped parenchyma cells, vascular bundles and stomata. The waxy cuticle layer reduces the rate of water loss from the surface of the leaf. The tough layer of epidermal cells helps in protection. The aerenchyma cells allow the transport of oxygen from leaves to the roots and play an important role in survival of rice plants. The stomata present on the upper epidermal cells are responsible for the water and gas exchange in and out of the system. The parenchyma cells are responsible for photosynthesis in plants. The vascular bundles help in transporting water and nutrients to various parts of the plants. All these structural observations are very much important, as they are involved in the survival and growth of plant.

The A-scan, B-scan and FESEM images for leaves under varied acidity levels provide the optical visualization of qualitative changes at microstructural levels. On the other hand, the scattering coefficient helps to quantify such changes. The analysis of A scans and B scans generated by SS-OCT system helped us to get an insight of the changes that take place in rice plant at tissue level under acidic stress. These images show progressive merging and disintegration of layers with increasing acidity. The FESEM images show reduction in stomatal density as well as reduction in stomata pore length and width with increasing acidity.

The increase in acidity is associated with decrease in photosynthesis, which can be explained by the alteration in leaves at microstructural level. The parenchyma cell layer associated with photosynthetic pigments starts disintegrating when acidity level is increased and finally merges with epidermal cell layer at very high acidity (pH 3.5). Our SS-OCT study shows these prominent changes occurring at different acidity levels in the layered organization of leaves in rice plants. In addition, calculation of scattering coefficient also shows distinct structural changes among leaves at pH 6.5, 5.5, 4.5 and 3.5. Lower scattering coefficient for leaves at pH 6.5 compared to other pH levels reflect the intact layered structure corresponding to healthy status.

The linkage between increase in acidity and reduction in photosynthesis can also be explained through morphological changes in leaves due to acidic stress. FESEM analysis reveals that higher acidity is associated with reduction in stomatal density and alteration in structure of stomata. The adverse changes in stomata characteristics at higher acidity leads to reduction in photosynthesis rate which is associated with stomatal conductance.

On the basis of results of SS-OCT and FESEM we can hypothesize that the microstructural alterations induced in leaves due to higher acidity are responsible for changes in physiological parameters. Thus, studying the microstructural changes at varied acidity levels will have a profound impact on agricultural science.

A detailed study on morphological characterization of rice leaf bulliform and aerenchyma regions by non-invasive and low coherence interferometry approach has also been carried out [55]. OCT has been applicable in detecting green mottle mosaic virus that is found in the seeds of cucumber. Two-dimensional OCT images and graphs revealed that the infected seeds had narrow gap between the seed coat and endosperm which was not present in the healthy seeds [75]. OCT has been more commonly utilized in different studies on animal model as compared to plant system. Recently OCT has been adopted to study microstructural changes in plant leaves under various conditions [51, 53-55]. However, the present work is the first report, where effect of low pH (acidity stress) on microstructural changes in rice leaves have been studied through SS-OCT technique.

The findings of our study with respect to FESEM analysis, which reveal that acidic stress have adverse effect on stomata characteristics leading to reduction in photosynthesis rate is similar to the effect of other stresses on stomata characteristics. A study on eggplants reported that ozone stress damaged stomatal structures and reduced the number of stomata. It was also found that stomatal density and pore size were also reduced. The rate of photosynthesis was reduced due to destruction of stomata which leads to reduced plant growth and lower yield [76].

## Conclusion

The present study reveals that acidic stress leads to the reduction in growth, as it negatively affects the plant’s height, chlorophyll content and various other growth parameters. The signs of stress and the response of plant were further corroborated through enhanced levels of TSS, proline content and peroxidised lipids. The results depicted that low pH inhibited growth of rice seedlings, which was caused by decreased chlorophyll content as well as reduced uptake of water. A relationship between acidic stress and microstructural changes at tissue level was also established using SS-OCT images and FESEM analysis. These microstructural changes were also associated with alteration of physiological parameters.

Our study is a holistic approach to understand the physiological, biochemical and microstructural changes occurring in the leaves of rice plant when subjected to acidic stress. In addition, we also hypothesize an inter-relationship between microstructural changes and physiological alterations. SS-OCT is very useful due to its greater penetration depth in high scattering samples like leaves. In agricultural studies, our approach can be applied to predict health of plants without affecting the yield of crops as only leaf sample is needed. The non-invasive SS-OCT based method reported in the study can be used to phenotype large population under acidic soil conditions. Overall this study represents the first SS-OCT based approach combined with FESEM analysis, along with various biochemical parameters such as TSS, proline content, lipid peroxidation, chlorophyll content and electrolyte leakage to study the effects of acidic stress treatment on rice plants. Our study thus charts out a new path to get detailed understanding of alterations in rice plant in response to acidic stress and the results of this study can be utilised to develop acidity tolerant rice cultivars in future.

## Acknowledgements

We thank Dr. Raju Poddar, Associate Professor, Department of Bio-Engineering, BIT Mesra, Ranchi, India for granting access to the Bio-Photonics Lab for this study. TEQIP–III fellowship given to Ms. Ekta is also acknowledged.

